# Variability in proliferative and migratory defects in Hirschsprung disease-associated *RET* pathogenic variants

**DOI:** 10.1101/2024.09.24.614825

**Authors:** Lauren E Fries, Sree Dharma, Aravinda Chakravarti, Sumantra Chatterjee

## Abstract

Despite the extensive genetic heterogeneity of Hirschsprung disease (HSCR; congenital colonic aganglionosis) 72% of patients harbor pathogenic variants in 10 genes that form a gene regulatory network (GRN) controlling the development of the enteric nervous system (ENS). Among these genes, the receptor tyrosine kinase gene RET is the most significant contributor, accounting for pathogenic variants in 12%-50% of patients depending on phenotype. RET plays a critical role in the proliferation and migration of ENS precursors, and defects in these processes lead to HSCR. However, despite the gene’s importance in HSCR, the functional consequences of RET pathogenic variants and their mechanism of disease remain poorly understood. To address this, we investigated the proliferative and migratory phenotypes in a RET-dependent neural crest-derived cell line harboring one of five missense (L56M, E178Q, Y791F, S922Y, F998L) or three nonsense (Y204X, R770X, Y981X) pathogenic heterozygous variants. Using a combination of cDNA-based and CRISPR-based PRIME editing coupled with quantitative proliferation and migration assays, we detected significant losses in cell proliferation and migration in three missense (E178Q, S922Y, F998L) and all nonsense variants. Our data suggests that the Y791F variant, whose pathogenicity has been debated, is likely not pathogenic. Importantly, the severity of migration loss did not consistently correlate with proliferation defects, and the phenotypic severity of nonsense variants was independent of their position within the RET protein. This study highlights the necessity and feasibility of targeted functional assays to accurately assess the pathogenicity of HSCR-associated variants, rather than relying solely on machine learning predictions, which could themselves be refined by incorporating such functional data.

## Introduction

The increasing use of whole exome (WES) and whole genome (WGS) sequencing for clinical diagnosis by identifying pathogenic variants in patients has identified hundreds of thousands of such variants for many rare diseases. When this approach results in the identification of a known pathogenic mutation in a gene associated with a phenotype, the benefit of this approach is without question. However, despite substantial progress in both sequencing technologies and the development of analytical tools for identifying pathogenic variants, the majority of variants identified in patients are neither pathogenic nor benign but of unknown significance (Fung et al., 2020; Neveling et al., 2013; Seo et al., 2020). This is owing to our inability to scale functional studies for assessing the pathogenicity of disease-associated variants for most genes, with some notable exceptions (Starita et al., 2018).

One such uncommon oligogenic disorder where multiple genes and their pathogenic variants have been identified by sequencing is Hirschsprung disease (HSCR)(Gui et al., 2017; Tang et al., 2018; Tilghman et al., 2019). HSCR is a developmental disorder of the enteric nervous system (ENS) characterized by the loss of proliferative and migratory enteric neural crest-derived cells (ENCDCs) of the developing (embryonic) colon, leading to aganglionosis of variable segments of the gastro-intestinal tract. HSCR is classified by the length of colonic aganglionosis into short segment (80% of cases), long segment (15% of cases), and total colonic aganglionosis (5% of cases) forms, arising from the extent of the developmental dysregulation (Badner and Chakravarti, 1990). Sequencing studies in HSCR have identified rare coding pathogenic alleles (PAs) in 24 genes, most frequently at *RET* and *EDNRB*, and common non-coding enhancer variants at *RET*, *NRG1*, and *SEMA3C/D*. Additionally, large copy number variants have been found at nine loci. In total, 72% of affected individuals of European ancestry harbor at least one PA in these 35 genes and loci, explaining 63% of the population attributable risk (Tilghman et al., 2019).

Despite the large genetic heterogeneity in HSCR, the receptor tyrosine kinase gene *RET* is *the* major contributor to the pathophysiology of Hirschsprung disease (HSCR). Approximately 12% of isolated cases and 35-50% of familial cases carry rare PAs in its coding exons (Gui et al., 2017; Tilghman et al., 2019; Tomuschat and Puri, 2015), including missense, nonsense, frame-shift, and small deletions/duplications. Notably, there is no association between the type of variant and the severity of the disease (So et al., 2011; Tomuschat and Puri, 2015), although in short-segment disease variants cluster in the intercellular domain (Kashuk et al., 2005). All of these studies suffer from the ascertainment bias of studying obviously pathogenic variants (nonsense variants and those at conserved residues). Recent, machine learning studies have suggested that many others also likely contribute to pathogenicity (Brandes et al., 2023; Cheng et al., 2023)

Current studies to address the role of disease-associated *RET* variants are limited in both cell lines and mouse models (Kjaer et al., 2010; Nakatani et al., 2020; Wang et al., 2019), particularly for investigating whether and how these variants disrupt cell proliferation and migration, the hallmarks of HSCR. The lack of functional studies in HSCR is partly owing to the challenges in accessing relevant cell lines for testing, the complexity of protocols for deriving and culturing ENS cells, and the poor *in vitro* migration of these cells (Fattahi et al., 2016). Although animal models are useful, they are expensive, may not precisely model the human developmental process and have low throughput (Nakatani et al., 2020). An alternative is to develop a scalable human cell line system for HSCR variant assessment and study genotype-phenotype correlations. Here, we utilize a neural crest-driven neuroblastoma line, CHP212, which expresses key ENCDC markers, including genes that are part of the *RET* gene regulatory network (GRN) disrupted in HSCR (Chatterjee and Chakravarti, 2019; Chatterjee et al., 2023; Chatterjee et al., 2016). In this system, we assessed the cell phenotypes of migration and proliferation in five missense (L56M, E178Q, Y791F, S922Y, F998L) and three nonsense (Y204X, R770X, Y981X) HSCR-associated *RET* variants (**Table 1**).

**Table 1:**
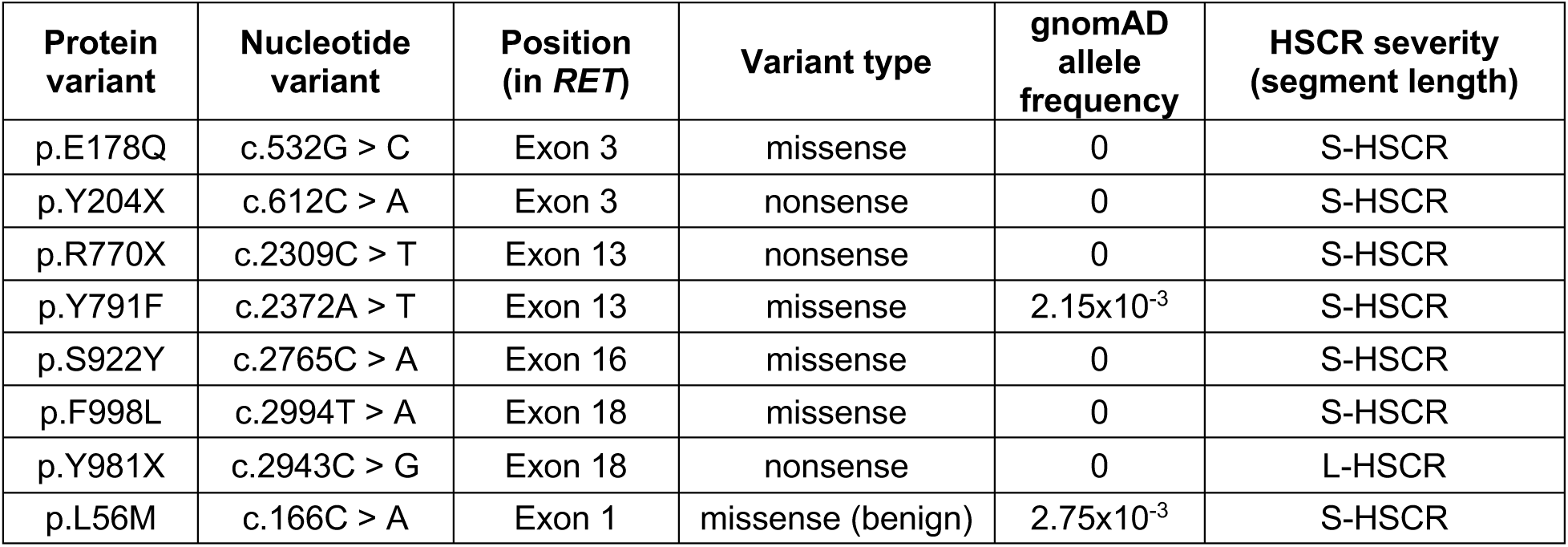
Nucleotide variants (ENSEMBL transcript: ENST00000355710.3) and the corresponding amino acid (UniProt id: P07949) changes in RET, and their gnomAD allele frequencies (v4.1.0), assayed in CHP212 cells for their cellular proliferation and migration phenotypes. The location of the RET exon containing a variant and the severity of the cognate patient’s HSCR are also shown.

| Protein variant | Nucleotide variant | Position (in *RET*) | Variant type | gnomAD allele frequency | HSCR severity (segment length) |
| --- | --- | --- | --- | --- | --- |
| p.E178Q | c.532G > C | Exon 3 | missense | 0 | S-HSCR |
| p.Y204X | c.612C > A | Exon 3 | nonsense | 0 | S-HSCR |
| p.R770X | c.2309C > T | Exon 13 | nonsense | 0 | S-HSCR |
| p.Y791F | c.2372A > T | Exon 13 | missense | $2.15 \times 10^{-3}$ | S-HSCR |
| p.S922Y | c.2765C > A | Exon 16 | missense | 0 | S-HSCR |
| p.F998L | c.2994T > A | Exon 18 | missense | 0 | S-HSCR |
| p.Y981X | c.2943C > G | Exon 18 | nonsense | 0 | L-HSCR |
| p.L56M | c.166C > A | Exon 1 | missense (benign) | $2.75 \times 10^{-3}$ | S-HSCR |

To test the effect of specific variants, we first generated a *RET* null CHP212 cell line in which we could express any individual mutant RET protein. This cell model thus allowed us to demonstrate that nonsense variants have a stronger effect on the migration of CHP212 *RET*-null cells as compared to missense variants, reflecting the underlying proliferative defect. This cell line, lacking endogenous RET, also enabled us to measure the effect of mutant proteins without the confounding influence of the wildtype endogenous protein. Additionally, we could also use CRISPR-based PRIME editing to engineer stable heterozygous variants, which we generated for one missense (Y791X) and two nonsense (Y204X, Y981X) tyrosine residues in RET for testing their proliferation and migration cell phenotypes. Our data provides evidence that variant type, rather than its position along the protein, correlates with disease severity. This finding offers a possible explanation for why a Y981X *RET* variant leads to long-segment HSCR (Emison et al., 2010) while a Y204X *RET* variant, despite being an early stop codon, only results in short-segment HSCR (Tilghman et al., 2019). This study now offers an approach and reagents for future testing of *RET* variants, and variants in other *RET* GRN genes, for assessing genotype-phenotype correlations.

## Results

### Creation of Ret depleted CHP212 line

To investigate the impact of *RET* disease-associated variants in HSCR, we utilized the neural crest cell-derived CHP212 neuroblastoma cell line. This line is particularly suitable for studying the differentiation of the neural crest lineage into the ENS due to its shared neural crest origin with *RET*-expressing ENCDCs *in vivo*. Gene expression analyses have confirmed the presence of key ENCDC markers in CHP212 cells (Harenza et al., 2017), including those within the *RET*-*EDNRB* GRN responsible for proliferation and migration of ENCDCs (Chatterjee and Chakravarti, 2019). To eliminate the confounding effects of any background *RET* variants its expression, we performed sanger sequencing of all *RET* exons in this cell line and found no nucleotide variants compared to the reference sequence. Additionally, we genotyped CHP212 for three HSCR-associated enhancer polymorphisms (rs2506030 (A/**G**), rs7069590 (C/**T**), rs2435357 (C/**T**); risk and expression reducing alleles in bold) to classify the cell line with respect to *RET* resistant (R: GTC, ATC, GCC, ACC) and susceptible (S: ATT, GTT) haplotypes. revious research demonstrated that the SS haplotype can significantly (4-fold) reduce *RET* expression in the human fetal gut, independent of coding variants (Chatterjee et al., 2023). Sequence analysis showed our cell line to be RR (ATC/ATC), so that coding variant effects can be quantified against a disease-resistant standard.

To assess the utility of CHP212 cells for cell migration studies, we generated a *RET*-depleted cell line via CRISPR-Cas9-mediated deletion employing a pair of guide RNAs targeting exon 2 of the *RET* gene. Sanger sequencing revealed an 18 bp homozygous deletion (chr10:43,100,456-43,100,473; hg38), encompassing part of the first intron and the first four amino acids of exon 2, present in 80% of the cells, with the remaining 20% being a mixture of heterozygous (16%) and wildtype (4%) cells. To evaluate the functional consequence of this deletion, we measured *RET* transcript and protein levels in these *RET*-deleted cells (*RET^del^*). We observed an 80% reduction in transcript levels (p<0.001) (**Figure 1A**) with no detectable RET protein by Western blot (**Figure 1B**). We next introduced a full-length *RET* reference sequence cDNA vector into *RET^del^*cells by transfection, restoring both transcript and protein levels to ∼90% of wild-type CHP212 levels (**Figure 1A, B**). To quantify the dependence of CHP212 cells on RET for cell migration, we performed scratch assays to measure the time required for wildtype and *RET^del^* cells to migrate across a gap: on average, *RET^del^* cells migrated 50% slower than wild-type cells. The addition of *RET* exogenously could partially rescue this phenotype, with migration rates reaching 90% of wild-type levels (**Figure 1C**). Taken together, these data demonstrate that the *RET^del^* cell line can serve as a good cellular model for assessing the functional consequences of *RET* HSCR-associated variants, when expressed from exogenous cDNA vectors, without the confounding effects of the endogenous protein.

**Figure 1:**
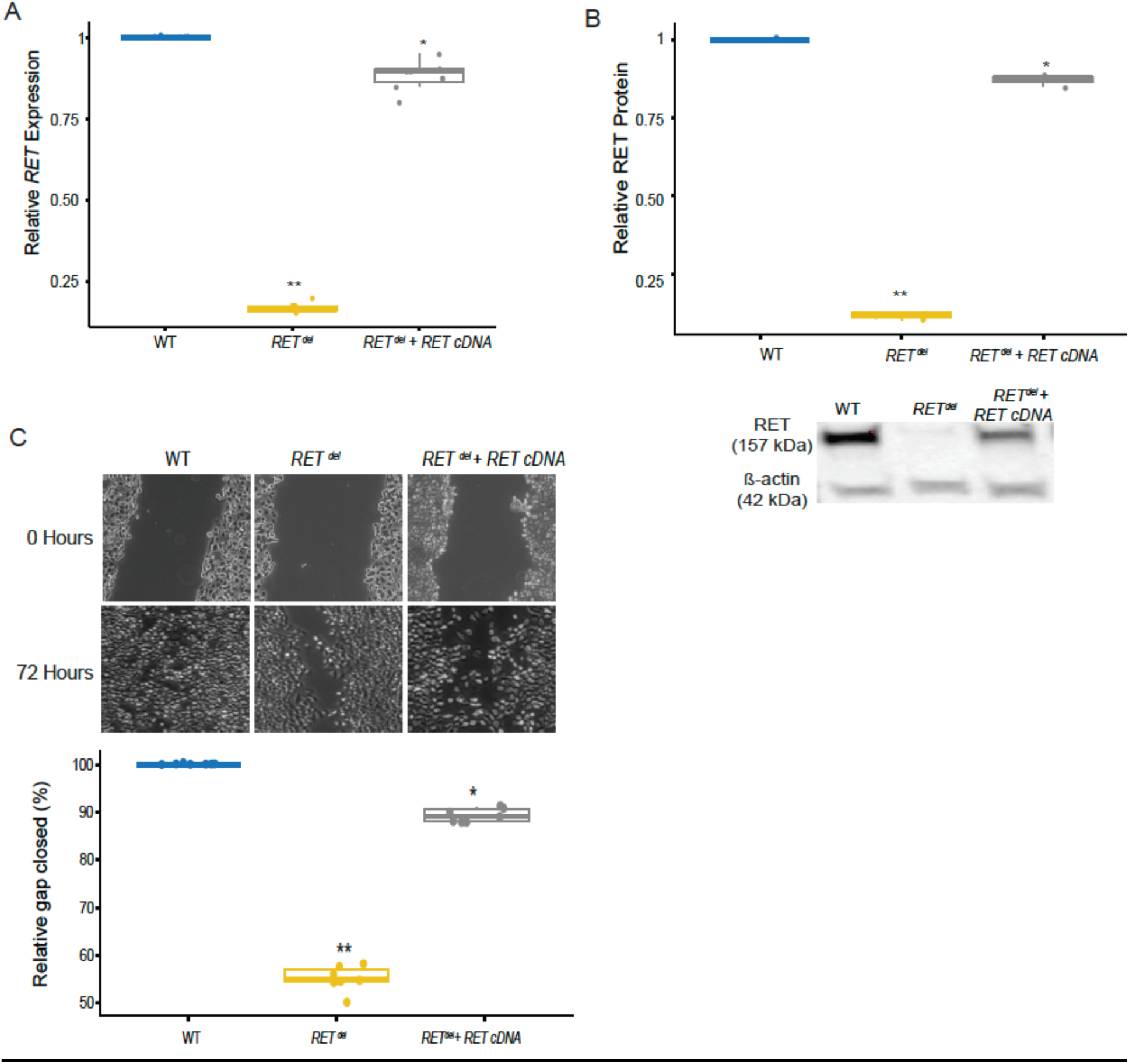
**(A)** CHP212 cells containing either heterozygous or homozygous *RET* deletions express only 10% *RET* transcripts compared to wildtype (WT), an effect which can be 90% rescued by expressing wildtype *RET* exogenously. **(B)** Concomitant changes can be observed in RET protein levels in the same cells. **(C)** Loss of RET function reduces cellular migration of CHP212 cells by 50% as compared to WT cells, which can be 90% rescued by addition of exogenous *RET* (* p<0.01, ** p<0.001).

### Migratory and proliferative defects in HSCR-associated RET coding variants

To test for the functional consequences of HSCR-associated *RET* variants, we focused on variants associated with the most common and low severity form of the disease - short segment HSCR in males with isolated aganglionosis. We choose 6 coding PAs with nonsense or missense changes in codons encoding amino acids at positions that are conserved across species. These variants were observed in our recent exome sequencing screen of 190 male White probands with S-HSCR at a frequency of 5% or less (Tilghman et al., 2019) (**Table 1**). Additionally, we also tested the Y981X nonsense variant which was detected by targeted *RET* sequencing of 2 probands with L-HSCR (Emison et al., 2010). As a negative control, we also selected a missense variant (L56M) previously reported as benign from functional tests to detect its effect on the production and folding of the RET protein (Kjaer et al., 2010) (**Table 1**).

To better understand the potential phenotypic consequences of the selected variants, we used AlphaMissense (Cheng et al., 2023) predictions to evaluate their pathogenicity (**Figure 2A**). Protein structure modeling by AlphaMissense predicted the E178Q, Y204X, R770X, S922Y and Y981X variants to be likely pathogenic. In contrast, Y791F and F998L were predicted as variants of uncertain significance (VUS), and L56M as likely benign (**Figure 2A**), consistent with prior functional tests (Kjaer et al., 2010). To experimentally test these predictions, we cloned full-length *RET* cDNAs, each containing one of the selected eight variants, into a mammalian expression vector with a high-efficiency CMV promoter. This vector also included an enhanced green fluorescent protein (EGFP) cDNA to monitor transfection efficiency. The mutant *RET* constructs were transfected separately into *RET^del^*cells, lacking endogenous RET expression, to introduce the mutant *RET* variants into a common genetic background. As a wildtype control, we transfected a wildtype *RET* cDNA into the same cell line. Thus, all *RET* expression measured, whether mutant or wildtype, originates from expression from the exogenously transfected vectors (**Figure 1A**).

**Figure 2:**
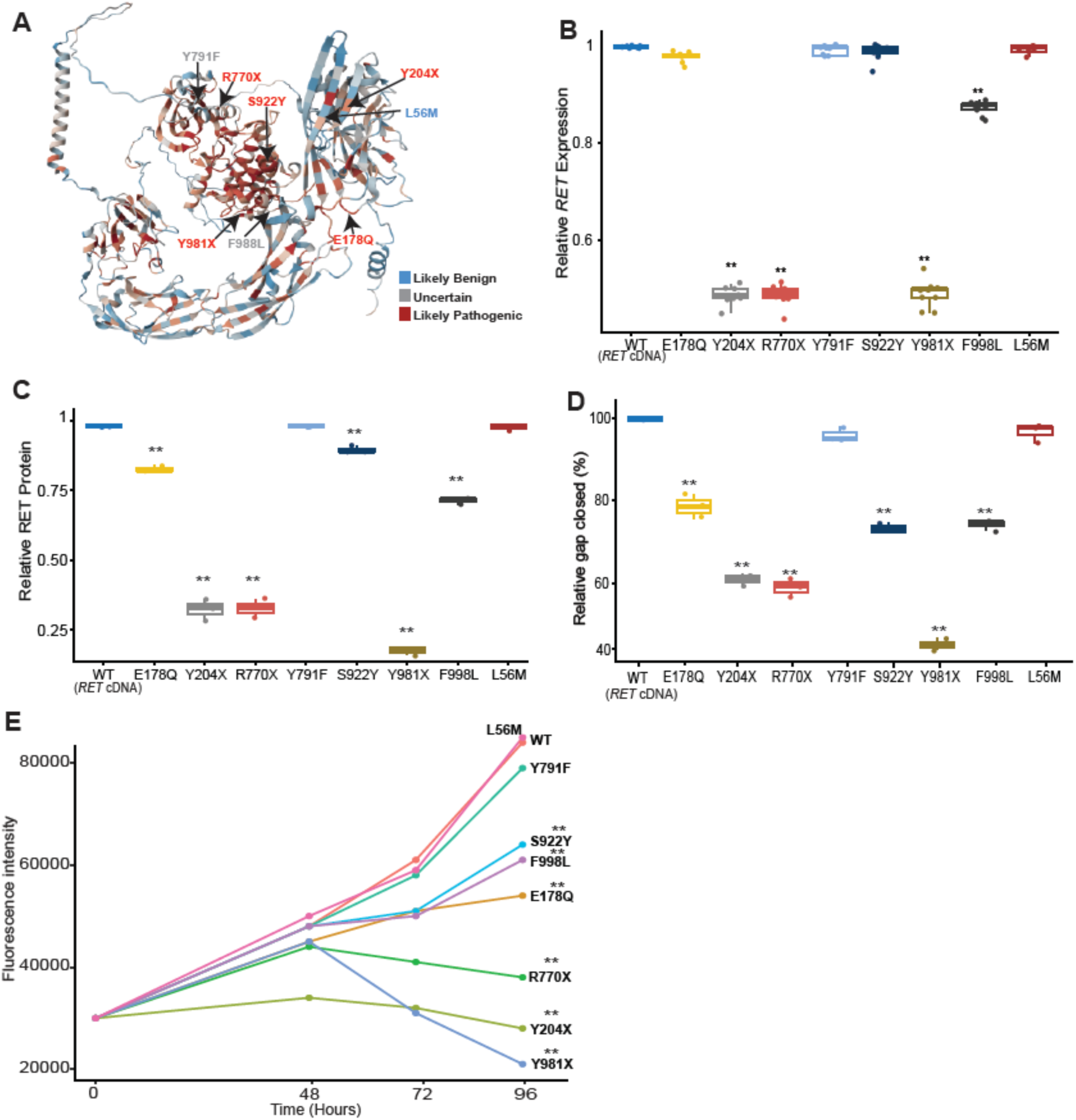
**(A)** 3-dimensional folding structure of full-length RET (UniProt id: P07949) as predicted by AlphaFold. All residues are colored by pathogenicity: red - likely pathogenic, blue - likely benign and grey - uncertain significance. **(B)** Expression of exogenously transfected RET protein containing HSCR specific variants. All nonsense variants express at 50% level as compared to wildtype (WT) RET but one missense variant (F998L) shows 15% reduced expression. **(C)** Relative fold change of each mutant RET protein as compared to the WT as measured by normalized density of bands from an immunoblot. **(D)** Scratch assays on all exogenously transfected mutant RET protein demonstrates significant migratory loss in all variants except Y971F and L56M. **(E)** All missense mutant cells proliferate normally till 48 hours but all nonsense variants (Y204X, R770X and Y981X) show significant loss of proliferation. At 72 and 96 hours all variants except L56M and Y791F show significant loss of proliferation as compared to WT cells, with nonsense variants show more significant loss than missense variants (* p<0.01, ** p<0.001).

We assessed the transfection efficiency of our individual vectors by quantifying EGFP expression using qPCR in the transfected cell lines, 72 hours post transfection. There were no significant differences in the normalized threshold cycle (ΔCt) values for EGFP, across different transfections (**Supplementary Figure 1**). Therefore, all subsequent measurements were assumed not to be influenced by variation in transfection efficiency.

We measured *RET* expression from individual mutant constructs at 72 hours post transfection. Among the constructs carrying missense variants, only F998L showed a 10% reduction in gene expression compared to wildtype (p=0.003), with no other mutants exhibiting any significant change (**Figure 2B**). This result is as expected, as mutations in the DNA typically do not affect transcript formation but may impact the resulting protein. The reduced expression in F998L may be due to the formation of unstable RNA secondary structures, a hypothesis that requires further investigation. Among the three nonsense mutants, Y204X showed a 46% reduction in *RET* gene expression (p<0.001), while R770X and Y981X exhibited reductions of 54% (p<0.001) and 52% (p<0.001), respectively (**Figure 2B**). The decrease in gene expression observed in these nonsense mutants is likely from nonsense-mediated decay (NMD), a genetic process which degrades mutant transcripts. NMD has been previously demonstrated to target transcripts produced from vectors in cell lines as well (Kim et al., 2017).

We next examined whether the changes in transcript levels we observed were reflected in RET protein output using Western blotting. Since RET is a cell surface receptor, we isolated membrane-associated proteins using membrane fractionation (Methods) and visualized them via immunoblotting 72 hours post-transfection, corresponding to the time point at which we could measure transcript changes (**Figure 2C, Supplementary Figure 2**). We used densitometric analysis to quantify protein levels. First, the missense variants, E178Q and S922Y, which showed no detectable change in mRNA levels, had 20% (p<0.001) and 15% (p<0.001) reductions in protein expression, respectively (**Figure 2C**). Second, F998L exhibited a 28% reduction in protein level, approximately three times greater than that observed at the transcript level (**Figure 2C**). This suggests that the F998L mutant protein undergoes additional degradation, which we hypothesize occurs by cellular mechanisms to clear misfolded proteins, although proof for this requires further study. Third, Y791F had no change in RET protein expression and neither did L56M, as expected from previous studies (Kjaer et al., 2010). Among the nonsense mutations, our antibody detected 30% of mutant protein for both Y204X and R770X and only 16% for Y981X (all three satisfied p<0.001) (**Figure 2C**). Given that NMD typically degrades transcripts containing nonsense mutations, we did not expect to observe any RET protein in these cells. Two possible explanations for this observation of residual protein are that either NMD is still active at 72 hours post transfection or that endogenous wildtype RET expression exists in a subset of cells where *RET* deletion failed. These results are from multiple replicates and so are robust to extreme replicate-to-replicate experimental variation.

We further investigated how such reduced RET expression could affect cell migration, a primary deficiency of ENCDCs in HSCR (Amiel et al., 2008; Montalva et al., 2023). We conducted scratch assays, as previously described, to measure the time required for wildtype and mutant cells to migrate across the gap generated by a scratch. After 72 hours, cells carrying the E178Q mutation had closed 80% of the gap as compared to wildtype cells (p<0.001); the S922Y and F998L mutants closed 72% (p<0.001) and 75% (p<0.001) of the gap, respectively (**Figure 2D**). In contrast, gap closure for the missense variants Y791F and L56M was near complete at 72 hours post-scratch, behaving like wildtype cells and reinforcing their classification as benign. Among the nonsense variants, Y204X and R770X had 60% (p<0.001) gap closure while Y981X had only closed 45% of the gap (p<0.001), highlighting the severity of the cellular defects of these mutants (**Figure 2D**).

In HSCR, loss of cell migration is a result of the loss of proliferative capacity of ENCDCs as we demonstrated in the developing mouse gut of a HSCR model (Vincent et al., 2023). Thus, to determine whether the observed migratory defects in mutant cells were associated with a loss of proliferative capacity, we subsequently performed a resazurin salt reduction assay. This fluorometric assay measures the metabolic activity of live cells as a proxy for proliferation. We recorded the fluorometric output at 48-, 72-, and 96-hours post-plating. Up to 48 hours post-plating, there were no significant changes in the proliferative capacity of any of the mutant cells as compared to wildtype cells (**Figure 2E**), however, all 3 cell lines carrying the nonsense variants showed significant proliferative defects. Cells carrying the Y204X mutant protein demonstrated a 30% reduced proliferation (p<0.001) as compared to wildtype cells, while cells carrying Y770X and Y981X mutations had on average 8% (p<0.01) and 6% (p<0.01) reduced proliferation, respectively (**Figure 2E**). However, at 72 hours, we observed significant reduction in proliferation and cell survival of 16% for both S922Y and F998L, and 18% for E178Q (p<0.001 for all) variants (**Figure 2E**). Among the nonsense variants, R770X exhibited a 33% reduction in proliferative capacity (p<0.001), while Y204X and Y981X showed reductions of 47% and 50%, respectively, compared to wildtype levels (p<0.001 for all) (**Figure 2E**). At 96 hours, the reduction in cell survival and proliferation of mutant cells became more pronounced. The proliferation of S922Y cells decreased by 23%, while F998L and E178Q cells showed reductions of 27% and 35%, respectively, compared to wildtype cells (p<0.001 for all) (**Figure 2E**). Even more substantial losses in proliferation were observed in cells carrying nonsense mutations: R770X, Y204X, and Y981X all exhibited decreases of 54%, 67%, and 75%, respectively (p<0.001 for all) (**Figure 2E**). These results indicate that the observed migratory defects in these cells are due to an underlying proliferative defect, leading to reduced cell survival, similar to what we observed *in vivo* (Vincent et al., 2023). Additionally, the variants Y791F and F998L were classified as variants of uncertain significance (VUS) by AlphaMissense and our assays suggest that Y791F is benign but F998L is pathogenic.

Taken together these data suggest that the nature of the protein impact, missense versus nonsense, rather than its position along the *RET* protein may determine the magnitude of its effect, i.e., that an early stop gain mutation in the protein doesn’t necessarily create a more severe phenotype. Our data also demonstrates that migratory defects observed in different disease-associated variants are intrinsically tied to cell survival and proliferation, i.e., reduction of the progenitor pool due to cell death leads to fewer viable cells which can migrate along the length of the gut.

### The effect of type and position of the variant on HSCR pathophysiology

Given our observation that nonsense variants, regardless of their position in the protein, more severely impaired *RET* function as compared to missense variants, led us to investigate this further. We utilized CRISPR-based PRIME editing to generate the specific nucleotide changes in stable CHP212 cell lines, each independently carrying either one of two nonsense variants (Y204X and Y981X) or the Y791F missense variant. Given RET’s role as a receptor tyrosine kinase, where multiple phosphorylated tyrosine residues are critical for signaling (Ibanez, 2013; Mulligan, 2014), we specifically targeted tyrosine residues throughout the protein (**Figure 3A**). We focused on the Y981 variant in the kinase domain which gets phosphorylated and Y204 present in the extracellular domain which has no reported phosphorylation (Encinas et al., 2004; Plaza-Menacho et al., 2016). Among the missense variants, we chose Y791F owing to its association with both multiple endocrine neoplasia, a gain-of-function (Plaza Menacho et al., 2005; Wagner et al., 2012), and HSCR, a loss-of-function (Seri et al., 1997; Tilghman et al., 2019) defect of RET.

**Figure 3:**
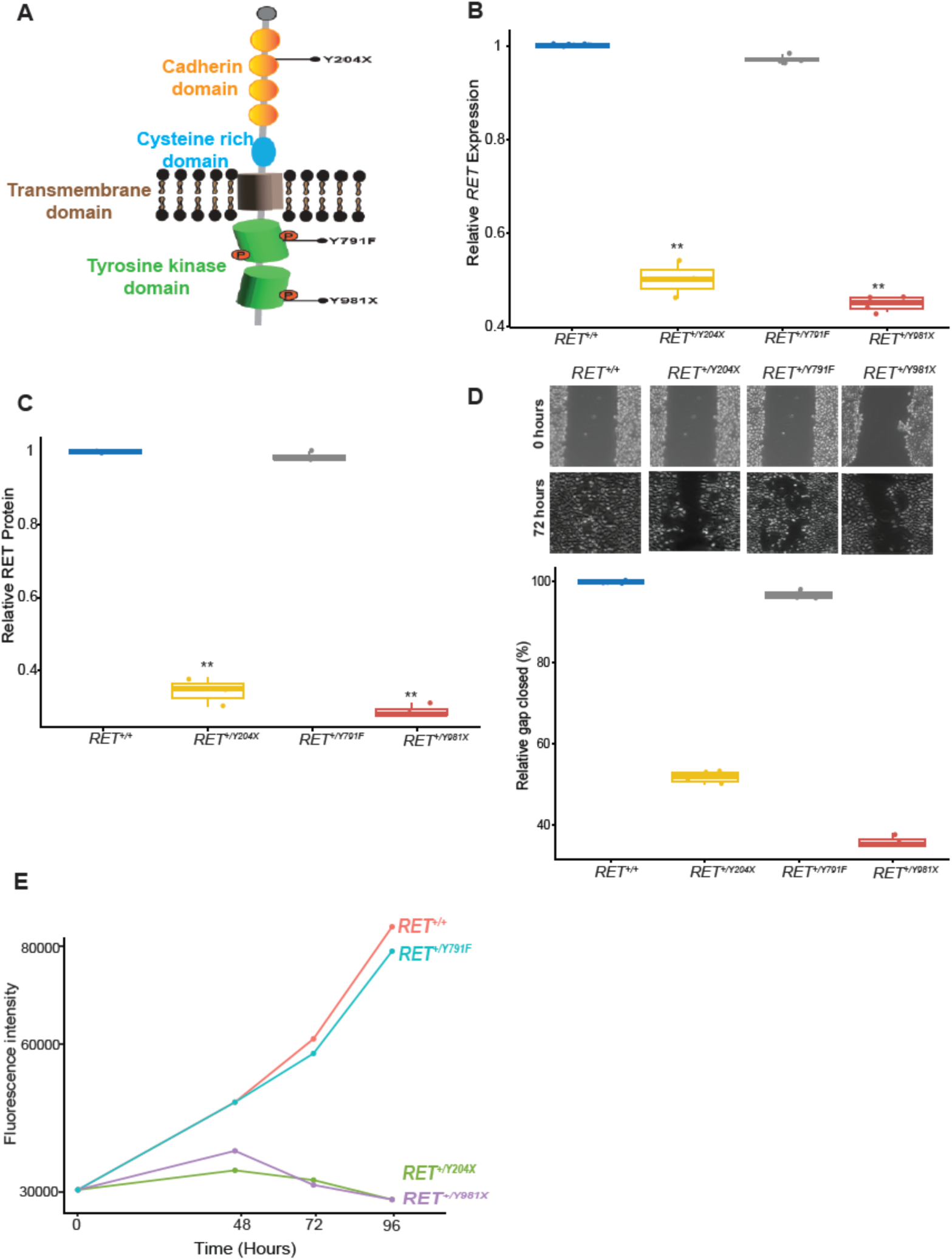
**(A)** Relative position of 3 tyrosine residues (Y204X, Y791F and Y981X) relative to the specific domains of the RET protein. **(B)** Loss of gene expression of *RET* observed in cell lines carrying heterozygous Y204X and Y981X variants. No expression change is observed in the Y791F variant cell line. **(C)** Loss of gene expression is reflected in the corresponding reduced RET protein in variant cell lines as quantified from an immunoblot. **(D)** Significant loss of migratory potential in *RET^+/Y204X^* and *RET^+/Y981X^* cells as compared to wildtype (WT) cells but with no effect in *RET^+/Y791F^* cells. **(E)** Rate of cell proliferation is severely reduced in both *RET^+/Y204X^* and *RET^+/Y981X^* cells 48 hours post plating with no cellular loss observed in *RET^+/Y791F^* cells as compared to WT cells.

The pathogenicity of Y791F has been debated: early studies reported a loss of cell migration upon introducing the mutation into HEK293 cells (Hyndman et al., 2013), however, more recent studies in cancer cell lines (Toledo et al., 2015) and the developing mouse ENS (Nakatani et al., 2020) found no effect of this variant, either *in vitro* or *in vivo*. Our cDNA-based studies of this variant also had no effect on migration and proliferation of CHP212 cells (**Figure 2E, 2F**); therefore, we wanted to determine if this lack of pathogenicity is observed in stable cell lines as well. We screened 48 single cell clones for all three mutants by Sanger sequencing and identified 4, 3 and 4 heterozygous clones for Y204X (*RET^+/Y204X^*), Y981X (*RET^+/Y981X^*) and Y791F (*RET^+/Y791F^*), respectively, suggesting an overall editing efficiency at the *RET* locus of 6-8%. We first measured *RET* transcript and protein expression in 2 stable lines for each variant by qPCR and Western blotting. In comparison to wildtype CHP212 cells (*RET^+/+^*^)^, both *RET^+/Y204X^* and *RET^+/Y981X^* had on average only 50% *RET* expression (**Figure 3B)**, although the corresponding protein levels were at 30% for both these heterozygous mutant cells (**Figure 3C and Supplementary Figure 3**). This observation of significantly lower protein levels than the expected 50% suggests that some of the wild-type protein may be degraded due to the presence of mutant protein, potentially suggesting that mutant RET protein can act in a dominant negative manner, an aspect that requires further investigation. There were no measurable changes in either transcript or protein levels of RET in *RET^+/Y791F^*cells as before (**Figure 3B, 3C**).

We finally performed scratch assays to determine if the migratory potential of these cells is compromised. At 72 hours post scratching, there were no significant differences between gap closure in *RET^+/+^* versus *RET^+/Y791F^* cells (p=0.16) (**Figure 3D**), but *RET^+/Y204X^*and *RET^+/Y981X^* cells closed only 50% (p<0.01) and 30% (p<0.01) of the gap, respectively, highlighting the severity of the cellular phenotype due to these nonsense mutations (**Figure 3D**). To also measure the corresponding changes in their proliferative capacity we again performed resazurin salt reduction assays. The *RET^+/Y791F^*cells continued to proliferate at the same rate as *RET^+/+^* cells up to 96 hours post plating further highlighting that this variant does not affect wildtype RET functions (**Figure 3E**). In contrast, *RET^+/Y204X^* cells have an decreasing 30%, 47% and 66% (p<0.01 for all) reduction in proliferation as compared to *RET^+/+^*at 48, 72 and 96 hours respectively; the corresponding decreasing reduction in proliferation for *RET^+/Y981X^* cells are 20%, 50% and 66% (p<0.01 for all) at the same time points, respectively. Thus, in our stable cell line assays, Y204X and Y981X cells show identical loss in their proliferative capacity at 72 and 96 hours, although the rate of migration is significantly (p=0.002) reduced in *RET^+/Y981X^* as compared to *RET^+/Y204X^*at 72 hours, as we observed in our prior scratch assays (**Figure 3D**). This uncoupling of migratory from proliferation defects suggest that there are additional yet undiscovered mechanisms which underlie the migratory defect in *RET^+/Y981X^*mutants and the severity of their migratory phenotype might explain why it led to L-HSCR (Emison et al., 2010) as compared to *RET^+/Y204X^*which was identified in a S-HSCR patient (Tilghman et al., 2019).

## Discussion

Just as the pace of gene and variant discovery have exploded owing to high throughput sequencing of genes and genomes, so has the need for functional characterization of disease-associated variants. We see two challenges for their solution. First, the relationship between a variant type (molecular change, position within its protein structure, frequency) and pathogenicity is poorly and incompletely understood. Moreover, this is likely to vary with a protein’s structure, e.g., extra-vs inter-cellular, exposed vs buried contacts, intra protein communication of signals. Second, in complex and polygenic disorders, whose molecular physiology at the level of genes lags far behind that of Mendelian disorders, pathogenicity presumably arises from its polygenic genotype, a biology that is even more poorly and incompletely understood. Therefore, we require cell models and a diverse array of cellular phenotypes for understanding how some genotypes ‘cause’ disease.

With regard to the first problem identified in the above, there has been a steady increase in studies performing experimental validation of disease associated variants to separate causality from association. These efforts have benefitted from high throughput testing of 1000s of variants associated with a particular gene using exogenous vectors (Findlay et al., 2018; Jia et al., 2021; Starita et al., 2018) or more focused approaches studying a few well-defined variants by genome engineering a specific gene to create stable cell lines (Shepherdson et al., 2024). The literature on scanning mutagenesis of individual genes is rapidly expanding and is the training set for deep learning and AI models to predict their pathogenicity (Brandes et al., 2023; Cheng et al., 2023), but these are not perfect tools. For example, for the RET variants we experimentally tested, AlphaMissense’s prediction matched 75% of variants. For the 2 variants predicted to be VUS, cellular assays demonstrated one to be benign (Y791F) and the other (F988L) pathogenic, indeed significantly affecting both proliferation and migration of cells. Consequently, improved deep learning will require much deeper cell assays on hundreds of variants of individual genes to increase their predictive power.

The second problem identified in the above is far more challenging. First, for polygenic disorders, where all the relevant genes are never specifically known, it is harder to establish pathogenicity which is a property of cells and tissues, not that for a single gene. However, we can make progress when single genes play an overwhelming role, as is the case with HSCR, where *RET* (and its interactor *EDNRB*) and its GRN explains 63% of disease attributable risk. An additional advantage is that the pathological HSCR phenotype is well defined, quantifiable and is largely restricted to a single cell type, ENCDCs (Vincent et al., 2023). The studies described herein are relevant because we have been able to test disease-associated variants in a disease relevant cell type (neural crest derived cells) and tested relevant cell phenotypes (proliferation and migration). Our current study, demonstrates that the variant type, missense vs. nonsense, or a smaller rather than a larger decrease in *RET* protein levels, has a greater impact on disease phenotype severity than the specific position of the mutation within RET. With respect to their effects on cells, this is exemplified by the similar proliferative defects but differing migratory impairments of Y204X (extracellular) and Y981X (intracellular) variants. The severe migratory defect of Y981X could explain its association with L-HSCR, highlighting the importance of considering both type and functional consequences of mutations in genotype-phenotype correlations.

In order to better understand HSCR, our challenge is to generate these cell assay data on all discovered HSCR-associated variants, rare or common, and other conceivable cell assays (e.g., receptor-ligand interactions), for deep learning of HSCR pathogenicity by genotype. At least for HSCR, we have identified patients with multiple pathogenic variants in multiple genes across the *RET-EDNRB* GRN (Tilghman et al., 2019): these are now important to study. Thus, we can both examine how different variants of specific genes affect the proliferation and migration phenotypes, as we have done here, but also how different genes within the GRN contribute to these two phenotypes. Further development of these assays for high throughput analyses are necessary both for screening the effects of newly discovered genes as well as to test the effect of targeted therapies, thereby offering powerful tools for dissecting the molecular mechanisms underlying HSCR.

## Materials & Methods

### CRISPR-Ca9 induced RET deletion

The second exon of RET was targeted using two guide RNAs: RETg1 (GUAGAGGCCCAAUGCCACUG; cut site: chr10: 43,100,459; GRCh38/hg38) and RETg2 (AAGCAUCCCUCGAGAAGUAG; cut site: chr10: 43,100,475 GRCh38/hg38) by transfecting a ribonucleoprotein complex containing 100 pmol of specific gRNA, coupled with 5 µg/µL of TrueCut Cas9 nuclease (Thermo Fisher Scientific) in Lipofectamine CRISPRMAX solution (Thermo Fisher Scientific). Three wells containing approximately 150,000 CHP212 cells were independently transfected. To increase the efficiency of deletion, we re-transfected the cells with the same ribonucleoprotein mix a second time after 72 h. The cells were further grown for 48 h and then equally split into two tubes for DNA and RNA extraction. To confirm disruption of the *RET* gene, specific primers (RET_E2F: CTTGCTGTACGTCCATGCC and RET_E2R: CCAGTTGTTCTCATGCAGCC) were used to verify deletions by PCR followed by Sanger sequencing. We used the ICE tool (Conant et al., 2022) to estimate the percentage of cells carrying various insertion/deletions (INDELs).

### Plasmid cloning for PRIME Editing

Cloning of pegRNA plasmids and secondary nicking gRNA plasmids were performed according to previously described protocols (Anzalone et al., 2019). Briefly, oligos were created using http://pegfinder.sidichenlab.org/ and synthesized by Integrated DNA technologies. Sequences are available in the Supplementary Table 1. The pU6-pegRNA-GG-Vector was purchased from Addgene (cat# 132777) and digested with Bsa1-HFv2 (NEB cat# R3733) at 37℃ for 1 hour. A 2.2 kb fragment from the plasmid containing the origin of replication, U6 promoter, U6 poly-T termination sequence, and AmpR gene were isolated with the QIAquick gel extraction kit (Qiagen cat# 28704) followed by QIAquick PCR cleanup (Qiagen cat# 28104). After annealing the protospacer, extension, and scaffold oligos, the oligos were cloned into the pegRNA plasmid by Golden Gate Assembly, combining digested plasmid, annealed oligos, Bsa1-HFv2, and T4 DNA ligase (NEB cat# M0202): these sat at room temperature for 10 minutes, were then incubated for 15 min at 37 °C, 15 min at 80 °C, and finally held at 12 °C. The assembly reaction was then transformed using One Shot Top10 Chemically Competent Cells (Thermo cat# C404010) and the resulting colonies were sequenced by Genewiz using U6 universal primer. Secondary nicking gRNAs were cloned into lentiGuide-Puro plasmid (Addgene cat# 52963). After digesting the lentiGuide-Puro plasmid with BsmBI-v2 the larger band on the gel was purified using Qiaquick Gel Extraction Kit. The guide sequence oligos were annealed and phosphorylated using T4 PNK (NEB cat# M0201) and then the oligos were cloned into the digested lentiGuide-Puro plasmid using Quick Ligase (NEB cat# M2200S). This assembly was then transformed using One Shot Stbl3 Chemically Competent *E. coli* (Thermo Fisher Scientific cat# C737303) and colonies were sequenced by using universal U6 primer.

### Cell culture for CHP212 lines

CHP-212 cells were purchased from ATCC (cat# CRL-2273) and cultured in 1:1 EMEM:Ham’s F12 (ATCC cat# 30-2003; Thermo Fisher Scientific cat# 1176505), 10% FBS, and 1% Penicillin Streptomycin (Thermo Fisher Scientific), and incubated at 37 °C with 5% O_2_. They were passaged every 2 days using 0.05% trypsin (Thermo Fisher Scientific).

### Stable cell line generation

250,000 CHP-212 cells were plated into each well of a 6 well tissue culture-treated plate 24 hours prior to transfection. On the day of the transfection, for each well, 75 µL Opti-mem (Thermo Fisher Scientific), 1.5 µg of pCMV-PE2 (Addgene cat# 132775), 500 ng of cloned pU6-pegRNA-GG-Vector, 200 ng of cloned lentiGuide-Puro, and 4.5 µL of Neuromag (VWR cat# 103258-382) were combined and incubated at room temperature for 20 minutes. The Neuromag/DNA complex was added to the plated cells, the plate was set on top of the magnetic plate, and then incubated at room temperature. After 20 minutes the media was removed and replaced with fresh media and the cells were cultivated at 37°C in a CO_2_ incubator for 48 hours. The cells were trypsinized, spun down, and resuspended in Flow Buffer consisting of HBSS, 10% FBS, and 2.5% HEPES. Single cells were plated via FACs machine into a 96 well plate coated with Poly-D-Lysine at 50 µg/mL. After growing up the single cells, colonies were sequenced by Genewiz for the appropriate mutation. Y204X colonies were sequenced using primers for exon 3 of *RET*, Y791F colonies were sequenced using primers for exon 13 of *RET*, and Y981X colonies were sequenced using primers for exon 18 of *RET* (Supplementary Table 1) using Q5 2x Master Mix.

### Gene expression Taqman assays

Total RNA was extracted from CHP212 cells using TRIzol (Life Technologies, USA) and cleaned on RNeasy columns (QIAGEN, USA). 500μg of total RNA was converted to cDNA using SuperScript IV reverse transcriptase (Life Technologies, USA) using Oligo-dT primers. The diluted (1/5) total cDNA was subjected to Taqman gene expression (Thermo Fisher Scientific) using transcript-specific probes and primers for *RET* (Mm00436305_m1; Thermo Fisher Scientific). Human β-actin was used as an internal loading control, as appropriate, for normalization. Three independent wells for CHP212 cells were used for RNA extraction and each assay performed in triplicate (n = 9).

The mean of the ΔC_t_ values (C_t_ Condition - C_t_ Actin) for each replicate for each condition was calculated after converting them to a linear scale (2^ΔCt^). The fold change was calculated as the ratio of the mean of 2^ΔCt^ experiment / 2^ΔCt^ control. The mean for the control was set to unity and the relative fold change was calculated based on these values. For calculating the variance of the of the fold change we used the following formula: 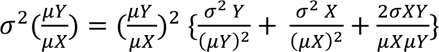 where *σ*^2^ is variance, *μ* is mean and 2*σXY* is the covariance between an experimental condition Y and its corresponding control X. Subsequently, P-values were calculated from pairwise 2-tailed t tests, and the data presented as the fold change with its standard error.

### Protein isolation and western blotting

10 cm plates of 90% confluent cells were trypsinized, spun down, and washed 3 times with PBS. Membrane protein was then extracted using the Mem-PER Plus Membrane Protein Extraction Kit (Thermo Fisher Scientific cat# 89842) as per the manufacturer’s instructions. The protein concentration was measured using Pierce BCA Protein Kit (Thermo Fisher Scientific cat# 23227) according to the manufacturer’s protocol. SDS-Page was performed by running 40 µg of protein on Bolt 4-12% Bis-Tris-Plus gel (Invitrogen cat# NW04120BOX). After washing the gel in 10% EtOH for 10 minutes, the iBlot Dry Blotting System (Thermo Fisher Scientific) was used to transfer proteins to a nitrocellulose membrane, using a program of 20 V for 2 m, 23 V for 5 m, then 25 V for 3 m. After washing the blot in PBST (1% Tween-20 in PBS) for 5 minutes 3 times, the blot was incubated overnight with shaking at 4°C in Blocking Buffer (PBST with 4% milk powder). The next day the blot was washed 3 times for 5 minutes in PBST before adding 1:500 dilution of anti-Ret antibody (Abcam: ab134100) in PBST and with overnight shaking at 4°C. The next morning 1:2,500 dilution of an anti-beta Actin antibody (Abcam: ab8227) was added to the blot and left with shaking at room temperature for 1 hour. After washing with PBST 3 times for 5 minutes the blot was incubated with Goat anti-Rabbit IgG (H+L) Secondary Antibody HRP (Thermo Fisher Scientific cat# 31460) at room temperature with shaking for 1 hour. The blot was washed for 5 minutes 3 times with PBST and then Pierce™ ECL Western Blotting Substrate (Thermo Fisher Scientific cat# 32106) was added, and the blot was imaged following the manufacturer’s instructions. To quantify the level of expression of each protein, we used ImageJ to measure the density of each band normalized to its corresponding loading control (β-actin). Relative fold change was measured compared to normalized density of the wildtype protein.

### Scratch assays

2.3×10^6^ CHP212 cells were plated in each well of a 6 well plate at T-1 in conditioning media, if applicable. 24 hours later a scratch was made across each well by hand with a P20 pipette tip. The wells were washed and then new normal or conditioning media was added. An image was taken at 10X as the T0 measure. 24 hours later the cells were gently washed, the media was replaced, and images was taken at 72 hours post scratching. The area of the scratch was quantified using PyScratch (Garcia-Fossa et al., 2020). The amount of gap closed by the wildtype cells at 72 hours post scratching was set to100%.

### Proliferation assays

50,000 CHP212 cells were plated per well in opaque-walled 96-well plates containing a final volume of 100 µl/well of DMEM/F12 media (Thermo Fisher Scientific). At 2 hours post plating, 20 µl resazurin solution was added to each well. This was considered to be time 0 for the experiment. Resazurin solution was added to 3 additional wells with no cells and only media, to serve as the background control. Cells were incubated with the solution for 4 hours and fluorescence was measured using a 560 nm excitation / 590 nm emission filter set. Similar measurements were repeated at 48, 72 and 96 hours. The data are presented as the mean fluorescent intensity for 3 wells per time point per variant.

## Author Contributions

AC and SC conceived and designed the study. LEF created all constructs, performed genome engineering experiments and cell proliferation and migration assays. SD conducted cell proliferation and migration assays; LEF, AC and SC wrote the manuscript. All authors approved of the final version of the manuscript.

The authors declare no conflict of interests.

