## Supplemental Figures and Tables for "Variability in proliferative and migratory defects in Hirschsprung disease-associated *RET* pathogenic variants"

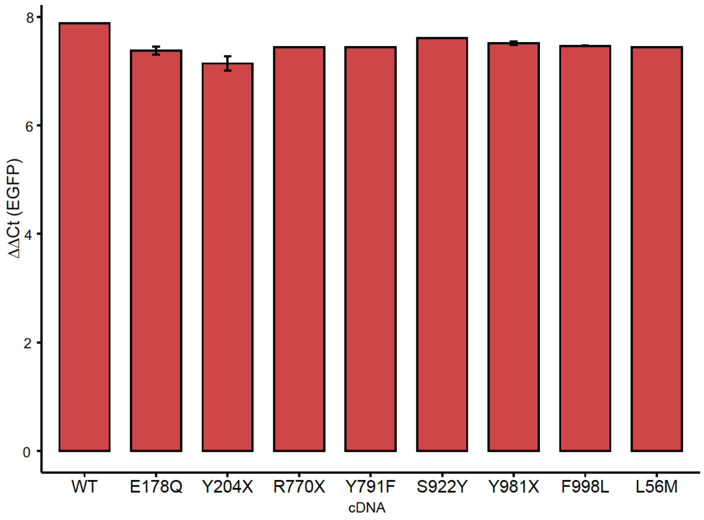


**Supplementary Figure 1**: Expression levels (Threshold cycle (Ct) values) of enhanced green fluorescent protein (EGFP) from individual overexpression vectors containing wildtype and mutant *RET* cDNA. There is no significant difference in EGFP expression between vectors highlighting equal transfection efficiency of each vector

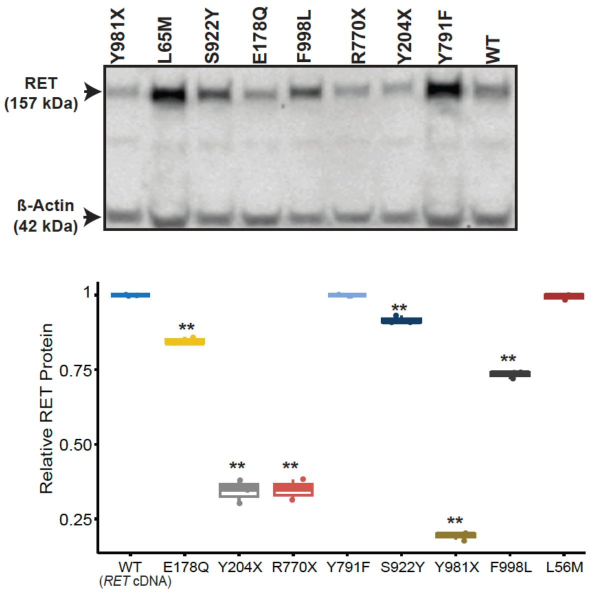


**Supplementary Figure 2**: Immunoblotting detects reduced RET protein for all nonsense variants and the missense F998L from proteins expressed from cDNA vectors

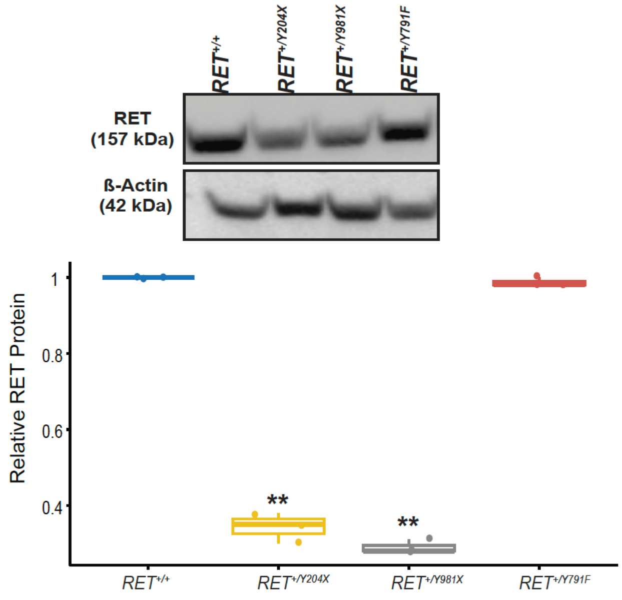


**Supplementary Figure 2**: Immunoblotting detects 30% wildtype RET protein in cells carrying the *RET*$^{+/Y204X}$ and *RET*$^{+/Y981X}$ mutations but no detectable change in RET protein levels for cells carrying the *RET*$^{+/Y791F}$ mutations

| Name | Sequence |
| --- | --- |
| Y204X pegRNA | CTCACCCTCCAGGAGCCTGTGTTTTAGAGCTAGAAATAGCAAGTTAAAATAAGGCTAGTCCGTTATCAACTTGAAAAAGTGGCACCGAGTCGGTGCAGCGTGGCCTAAAGGCTCCTGGAG |
| Y204X nicking RNA | caccgCAGCGTGGCCTAAAGGCTCC |
| Y791F pegRNA | caccgACATGTCATCAAATTGTATGgttttagaGCTAGAAATAGCAAGTTAAAATAAGGCTAGTCCGTTATCAACTTGAAAAAGTGGCACCGAGTCGgtgcGCTGCAGGCCCCAAACAATTTGATGACATGT |
| Y204X nicking RNA | caccgCAGGTCTCGCAGCTCACTCG |
| Y981X pegRNA | TCCAGCATTGCAGCATCAGGgttttagaGCTAGAAATAGCAAGTTAAAATAAGGCTAGTCCGTTATCAACTTGAAAAAGTGGCACCGAGTCGgtgcAGGTAGCGCCTGATGCTGCAATGC |
| Y204X nicking RNA | caccgCAAAGACCTGGAGAAGATGA |
| Exon 3 of RET F Sequencing Primer | AGCTCCTGCCTCCTCCCCATTC |
| Exon 3 of RET R Sequencing Primer | ATGGCTTGTGTCAAGGGCTCGC |
| Exon 13 of RET F Sequencing Primer | GTGGTTGCTGGCTCCTCA |
| Exon 13 of RET R Sequencing Primer | AGCGCGGGGCCCCTCTGATG |
| Exon 18 of RET F Sequencing Primer | AGCTCCTGCCTCCTCCCCATTC |
| Exon 18 of RET R Sequencing Primer | CACACTGGGAACTCTGAGG |

**Supplementary Table 1**: Sequences of PEG and nicking RNAs used to generated specific nucleotide changes in the *RET* gene to create stop codon at instead of tyrosine (Y) residues at amino acids 204 and 981 and convert tyrosine to phenylalanine at amino acid 791. Sequencing primers to detect these nucleotide changes are also provided for Exon 3 (c.612C > A; Y204), Exon 13 (c.2372A > T; Y791F) and Exon 18 (c.2943C > G; Y981X)
